# Glycosylation-dependent Turnover of Triterpenoid Saponins Controls Insect Deterrence

**DOI:** 10.64898/2026.05.04.721528

**Authors:** Jincheng Shen, Pablo D. Cárdenas, Søren Bak

## Abstract

**Background and Aims:** Plants deploy triterpenoid saponins as chemical defences against herbivores, yet it remains unclear whether insect digestion detoxifies these compounds or generates equally or more active metabolites. Because saponin bioactivity depends strongly on glycosylation patterns, we examined the fate and defensive activity of hederagenin-derived saponins during herbivory.

**Methods:** Larvae of *Plutella xylostella* were fed leaf discs containing structurally defined hederagenin-derived saponins. Saponin composition in treated leaves and larval frass was analysed by LC– qTOF–ESI–MS/MS. Feeding assays were used to compare the antifeedant activity of mono- and bidesmosidic forms.

**Key Results:** Larvae selectively metabolized complex hederagenin-derived saponins into simpler forms, with cellobiosides converted into monoglucosides during digestion, resulting in a marked shift in saponin composition between ingested material and frass. Feeding assays showed that monodesmosidic saponins strongly deterrer feeding, whereas bidesmosidic saponins were largely inactive. The loss of activity in bidesmosidic saponins was not explained by differential metabolism, indicating that glycosylation patterns directly determine biological function.

**Conclusions:** Insect herbivores selectively modify saponin structures through deglycosylation, thereby altering their defensive properties. Our findings demonstrate that glycosylation governs both saponin activity and metabolic fate, highlighting insect-driven turnover as a critical component of plant chemical defence during plant–herbivore interactions.

**Issue Section:** Original article

## INTRODUCTION

Plants produce a vast repertoire of specialized metabolites that function as chemical defences against herbivorous insects, shaped by long-term coevolutionary interactions (Divekar et al., 2022; Fürstenberg-Hägg et al., 2013). The diversification of these compounds has been facilitated by gene duplication and neofunctionalization, generating extensive structural and functional variation that enhances defensive capacity (Qiao et al., 2019). Among these metabolites, triterpenoid saponins are widespread defence compounds that cause toxicity and antifeedant effects against herbivores in a structure-dependent manner (Osbourn, 2003; Augustin et al., 2011; Moses et al., 2014; Cárdenas et al 2019). Glycosylated oleanolic acid- and hederagenin-derived saponins are prominent examples in which biological activity is strongly influenced by glycosylation pattern (Liu et al., 2019; Tian et al., 2021). Variation in sugar number, identity, linkage, and position of sugar moieties plays a central role in modulating their physicochemical properties and biological effects (Gao et al., 2011; Tian et al., 2021). However, the precise structural features that determine their activity in plant– insect interactions remain incompletely understood.

Glycosylation not only diversifies plant defence compounds but can also regulate their activity, for example by facilitating storage, transport, and controlled activation while preventing autotoxicity. In parallel, insect herbivores have evolved diverse strategies to cope with plant chemical defences, including detoxification, tolerance, and sequestration. In several systems, specialist insects sequester glycosylated plant metabolites and repurpose them for their own defence, illustrating the close interplay between plant biosynthesis and insect metabolism (Morant et al., 2008; Fürstenberg; Pentzold 2014; Beran et al., 2019). A prevailing assumption is that insect herbivores detoxify glycosylated defence compounds through enzymatic hydrolysis, for example by β-glucosidases, thereby reducing their toxicity (Morant et al., 2008). However, this idea has rarely been tested directly for saponins, whose biological activity is closely linked to their amphiphilic structure and glycosylation state (Devishi et al., 2026; Augustin et al., 2011).

Studies of *Barbarea vulgaris* and its specialist herbivores have provided intriguing but inconclusive insights. Nielsen et al. (2010) showed that both hederagenin cellobioside and α-hederin strongly deterred susceptible flea beetles, whereas only α-hederin remained active in tolerant lines. This led to the hypothesis that tolerant beetles may be detoxify saponins via β-glucosidase-mediated deglycosylation (Nielsen et al., 2010). Yet the enzymes involved, the metabolic products formed, and their biological activities remain unresolved (de Jong et al., 2009; de Jong and Nielsen 2010). Recent work challenges the assumption that deglycosylation necessarily leads to detoxification. Monoglucosides derived from hederagenin-based saponins retain strong antifeedant activity and can even display increased toxicity toward herbivores, including flea beetles and the brassica specialist *Plutella xylostella* (Augustin et al., 2012; Liu et al., 2019). These findings raise the possibility that insect-mediated turnover of saponins may generate more, rather than less, active compounds. Thus, whether insect digestion reduces or enhances saponin toxicity remains an open question.

Despite these advances, direct evidence linking insect metabolism of saponins to changes in their chemical composition and defensive activity is lacking. Here, we investigated saponin turnover and defensive activity using crude extracts of G-type *Barbarea vulgaris*, which contain hederagenin cellobioside and related saponins (Agerbirk et al., 2003; Kuzina et al., 2009; Khakimov et al., 2016; Liu et al., 2019), together with the purified compounds α-hederin and hederacoside C. These compounds differ in their glycosylation pattern, allowing us to disentangle the role of sugar composition and substitution at the C3 and C28 positions. We combined insect feeding assays with LC–qTOF–ESI–MS/MS metabolite profiling to determine how saponins are processed by *P. xylostella* larvae and to identify structural features that govern their deterrent and toxic effects. Specifically, we tested the hypothesis that insect digestion alters saponin composition through selective deglycosylation and that glycosylation patterns determine defensive activity.

## MATERIALS AND METHODS

### Plant, chemical compounds and insect materials

*Barbarea vulgaris* G-type and *Brassica napus* were grown in soil in a greenhouse at 19°C under a 16:8 h light-dark photoperiod. *Plutella xylostella* (diamondback moth) was reared at 20°C under a 16:8 light-dark photoperiod. *B. napus* leaf discs (16 mm diameters), treated with *B. vulgaris* G-type leaf crude saponin methanol extract, α-hederin (monodesmosidic saponin, Merck Sigma Aldrich) or hederacoside C (bidesmosidic saponin, Merck Sigma Aldrich), were used in this study for nonchoice insect feeding assay. First-instar larvae were used for mortality assay because of their higher sensitivity (Shinoda et al., 2002), whereas third-instar larvae were used for feeding and turnover assay because they consume leaf material more rapidly and are more tolerant (Vats et al., 2019).

### Saponin extraction

Saponins were extracted as described by Liu et al. (2019). In short, 1 g of *B. vulgaris* G-type leaf material was ground under liquid nitrogen and extracted with 3 mL 80% methanol (v/v) by 30 min sonification, followed by centrifugation at 16,000 g for 10 minutes. Supernatants were filtered through a 0.22-μm filter. An aliquot of 200 µL of the filtered extract was collected and used for LC-qTOF-ESI-MS/MS injection. The remaining extract was applied to *B. napus* leaf discs for insect feeding assays. *B. napus* leaf material was extracted with the same method and injected to LC-qTOF-ESI-MS/MS as negative control. Larval frass was collected with a brush, suspended in 0.3 mL 80% methanol, and extracted using the same procedure.

### Insect feeding assay

To investigate insect-mediated saponin turnover, four-fold diluted 80% methanolic extracts of Barbarea vulgaris G-type leaves were applied to Brassica napus leaf discs (16 mm diameter). Purified α-hederin and hederacoside C were dissolved individually in 80% methanol and applied separately at 6.6 nmol cm^−2^ by pipetting.

After treatment, the *B. napus* leaf discs were dried under airflow to evaporate the solvent prior to feeding. Ten treated *B. napus* leaf discs (16 mm diameters) were placed in each petri dish (9.4 × 1.6 cm). A smaller PETRI DISH (5.3 × 1.4 cm) containing water was placed in the centre to maintain humidity, and the larger dish was sealed with parafilm. Third-instar larvae of *Plutella xylostella* were starved for 3-4 h before transfer to the assay dishes. Frass was collected after 24 h of feeding and extracted in 80% methanol as described above. Frass from larvae fed methanol-treated leaf discs served as a negative control.

To determine whether saponin turnover resulted from insect digestion rather than residual plant enzymes (e.g. β-glucosidases), identically treated discs were incubated without larvae, then ground without liquid nitrogen to retain enzyme activity. Saponins were subsequently extracted as described above.

To evaluate the insect deterrence ability of α-hederin and hederacoside C, consumption area was quantified when approx. half of the *B. napus* leaves were consumed and quantified by using ImageJ (approx. 12 h).

To assess antifeedant activity, leaf area consumed from discs treated with α-hederin or hederacoside C was quantified using ImageJ when approximately 50% of the methanol-treated control discs had been consumed (approximately 12 h).

### Metabolite analysis

Saponin detection and data analysis were performed as previously described by Shen et al. (2026). Saponins were identified and quantified by LC-qTOF-ESI-MS/MS using DataAnalysis software (Bruker Daltonics). Authentic standards of hederagenin cellobioside, α-hederin and hederacoside C were used for compound identification and retention time confirmation. Monoglucosides were identified based as previously described (Augustin et al., 2012; Liu et al., 2019; Ertmann et al., 2018).

## RESULTS

### Insect digestion shifts saponin composition towards monoglucosides

To test whether G-type *Barbarea vulgaris* saponins are metabolized during insect digestion, crude methanolic extracts were applied to *Brassica napus* leaf discs, and the relative composition of hederagenin-derived saponins was quantified before and after ingestion by third-instar *Plutella xylostella* larvae using LC–qTOF–ESI–MS/MS.

Feeding by *P. xylostella* larvae resulted in a marked shift in saponin composition, indicating metabolic transformation during digestion. In plant extracts, hederagenin cellobioside, monoglucoside and other saponins accounted for 85%, 6% and 9% of the detected saponins, respectively (Fig. 2). Following ingestion, these proportions changed to 56%, 38% and 6% in larval frass, demonstrating a substantial enrichment of monoglucosides at the expense of hederagenin cellobioside (Fig. 2, Fig. 3).

**FIG. 1.**
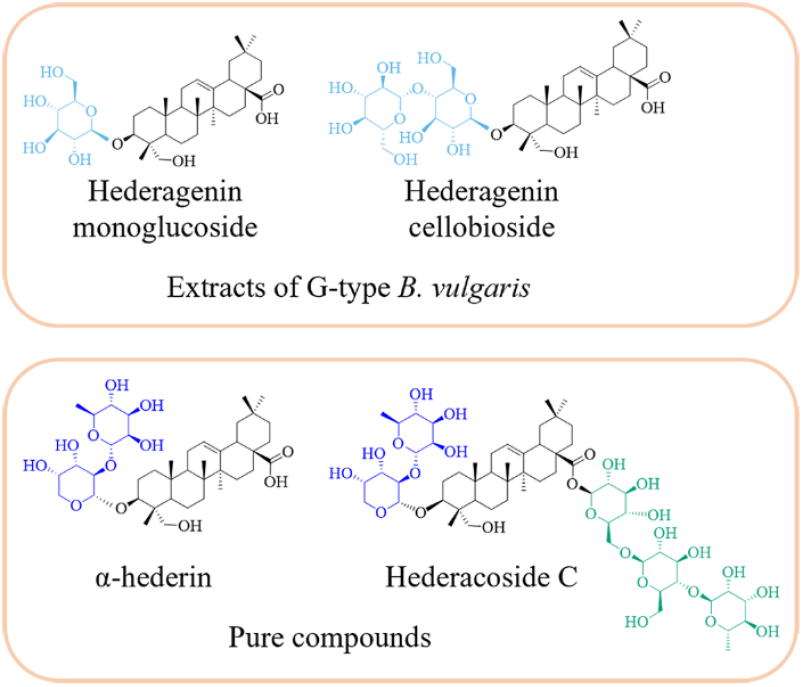
Triterpenoid saponins investigated in this study. The hederagenin aglycone is shown in black, while the sugar moieties are distinguished by different colors. All compounds share the hederagenin aglycone backbone. Monodesmosidic saponins: hederagenin monoglucoside (with one hexose) and hederagenin cellobioside (with an additional hexose at C3 compared to the monoglucoside), and α-hederin (with one rhamnose and arabinose). Bidesmosidic saponin: hederacoside C (with an extra sugar chain at C28 compared to α-hederin).

**FIG. 2.**
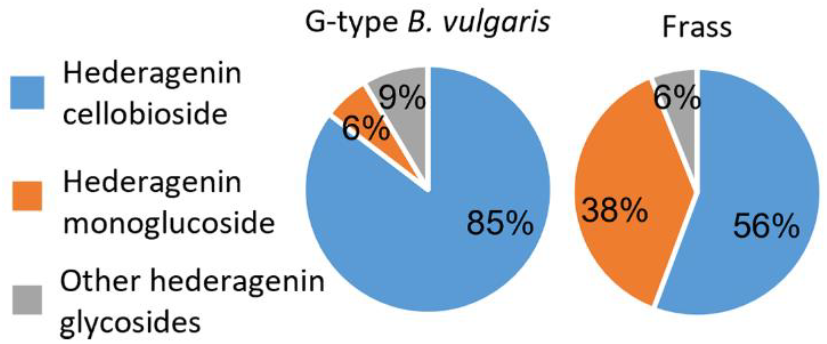
Insect digestion shifts hederagenin-derived saponins toward monoglucoside. Relative abundance of hederagenin-derived saponins in G-type *B. vulgaris* leaf extract and frass from *P. xylostella* larvae fed on the crude extract, as determined by LC-qTOF-ESI-MS/MS. Higher-glycosylated saponins were reduced during digestion, whereas the monoglucoside increased. Extracted ion chromatograms (EICs) for hederagenin-derived saponins: m/z = 679.4063 ± 0.2 (one hexose), EIC: m/z = 841.4591 ± 0.2 (two hexoses), EIC: m/z = 1003.5115 ± 0.2 (three hexoses).

**FIG. 3.**
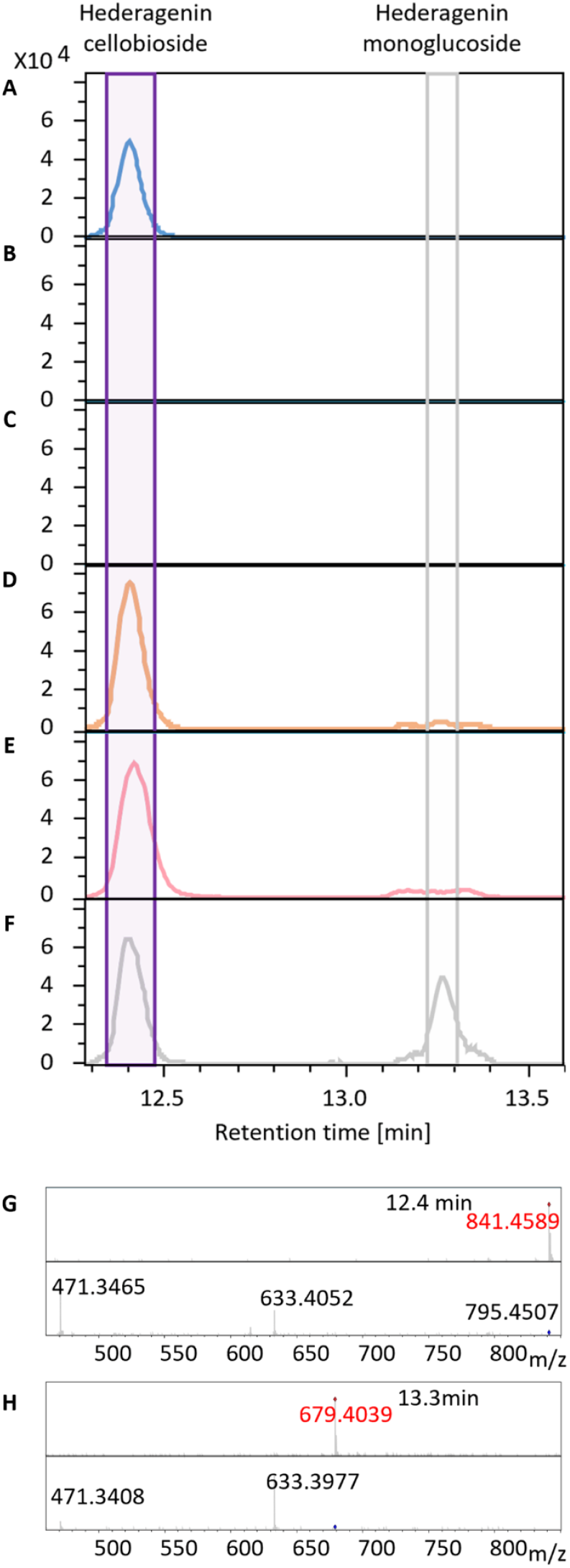
Hederagenin cellobioside is converted to monoglucoside during digestion by *P. xylostella* larvae. (A-F) Extracted ion chromatogram obtained by LC-qTOF-ESI-MS/MS for hederagenin cellobioside (EIC: m/z = 841.4597 ± 0.2, elucidated at 12.4 min) and hederagenin monoglucoside (EIC: m/z = 679.4063 ± 0.2, elucidated at 13.3 min). (A) Hederagenin cellobioside standard. (B) Extract from *B. napus*. (C) Extract from insect frass after feeding on *B. napus*. (D) Extract from the G-type *B. vulgaris*. (E) *B. napus* extract dipped with G-type *B. vulgaris* extract. (F) Frass extract from insects feeding on *B. napus*, dipped with G-type *B. vulgaris* extract. MS/MS fragmentation pattern of hederagenin cellobioside (G) and hederagenin monoglucoside (H) with two or one hexose units.

This compositional change is consistent with selective deglycosylation of hederagenin-cellobioside during insect digestion, generating the corresponding mono-glucoside. The apparent reduction of the other higher glycosylated saponins indicates that transformation is not restricted to hederagenin cellobioside (Fig. 2). Aglycones were not detected (data not shown), which suggest that hydrolysis did not proceed to complete deglycosylation under the conditions tested.

Control treatments, including *B. napus* leaf discs treated with *B. vulgaris* extract and incubated without larvae, as well as samples processed without liquid nitrogen, did not show this compositional shift (Fig. 3). These results indicate that the observed transformation resulted from insect digestion rather than plant enzymatic activity or sample handling. Because this process led to enrichment of monoglucosides, we next examined whether differences in glycosylation state influenced saponin biological activity.

### Glycosylation pattern determines antifeedant activity

To assess the functional consequences of glycosylation, we compared the antifeedant activity of structurally defined saponins differing in glycosylation patterns. α-Hederin (monodesmosidic, glycosylated at C3) and hederacoside C (bidesmosidic, glycosylated at both C3 and C28) share the same aglycone and identical C3 substitution but differ by the presence of an additional glycosyl chain at C28 (Fig. 4).

**FIG. 4.**
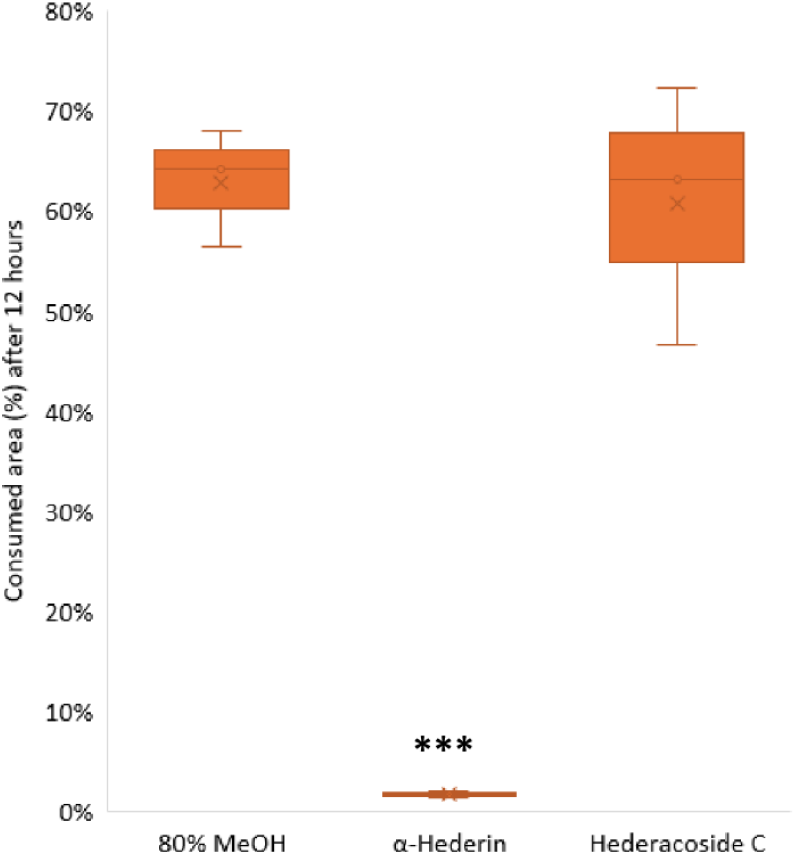
α-Hederin strongly reduces feeding by *P. xylostella* in a 12 h non-choice assay. Leaf area consumed (%) was measured when approximately 50% of methanol-treated control discs had been consumed. Data are presented as mean ± standard deviation. Statistical significance was assessed using Student’s t-test: *(p<0.5), ** (p < 0.05) and *** (p < 0.005).

Feeding assays revealed a striking difference in activity between these compounds. Leaf discs treated with α-hederin (6.6 nmol/cm^2^) were scarcely consumed, corresponding to an antifeedant effect of approximately 98%, whereas hederacoside C-treated discs (6.6 nmol/cm^2^) were consumed at levels comparable to the 80% methanol control (Fig. 4).

Metabolite analysis of larval frass showed that neither α-hederin nor hederacoside C was detected as lower-glycosylated products, indicating that these compounds were not measurably transformed under the experimental conditions. Thus, the observed difference in antifeedant activity cannot be explained by differential metabolism. Instead, the additional glycosylation at C28 abolished antifeedant activity. Given the strong deterrent effect of the monodesmosidic saponin α-hederin, we next tested whether this activity reflected direct toxicity.

### Monodesmosidic saponin α-hederin exhibit dose-dependent toxicity

To quantify toxicity, first-instar *P. xylostella* larvae were exposed to increasing concentrations of α-hederin in a non-choice feeding assay. Mortality increased in both a dose- and time-dependent manner. Dose–response relationships were analysed using a two-parameter log-logistic model (LL.2), allowing estimation of lethal doses (Dervishi et al., 2025). The corresponding EC_50_ values were 4.7 nmol/cm^2^ after 24 h and 3.0 nmol/cm^2^ after 52 h (Fig. 5), respectively, indicating both acute and delayed toxic effects. These results demonstrate that α-hederin is not only a strong feeding deterrent but also exerts direct toxic effects on insect larvae. Together with the absence of detectable turnover and strong antifeedant activity, this supports that monodesmosidic saponins, such as α-hederin, represent biologically active defensive forms, in contrast to the inactive bidesmosidic compound hederacoside C.

**FIG. 5.**
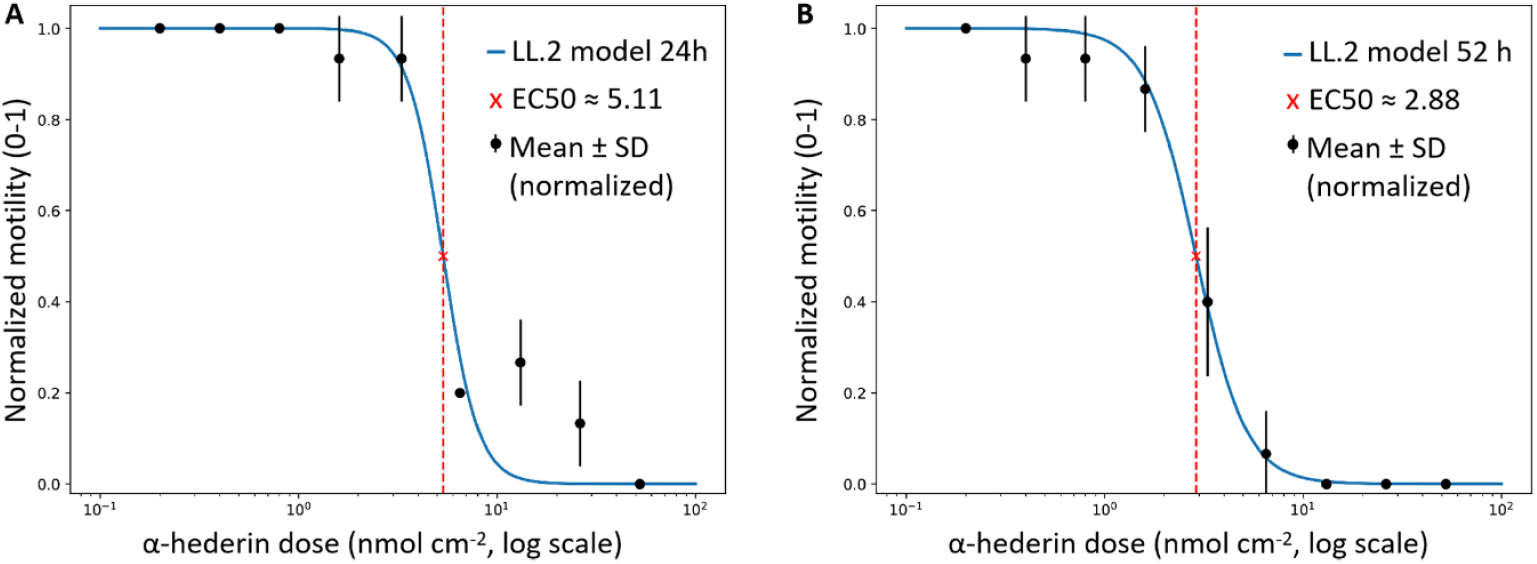
Dose-dependent mortality of first-instar *Plutella xylostella* larvae exposed to α-hederin. The EC_50_ of α-hederin for first-instar *P. xylostella* larvae was 5.11 and 2.88 nmol cm^−2^ after 24 h (A) and 52 h (B), respectively. Dose–response curves were fitted using two-parameter-log-logistic concentration-response model (LL.2) (Dervishi et al., 2026), which fixes the lower limit at 0 (no mortality in controls) and the upper limit at 1 (complete mortality at the highest concentrations).

## DISCUSSION

### Insect digestion drives selective partial deglycosylation of hederagenin-derived saponins

In this study, we show that digestion by *Plutella xylostella* larvae markedly alters saponin composition, leading to enrichment of monoglucosides at the expense of higher-glycosylated forms. The absence of detectable free aglycones indicates that hydrolysis does not proceed to complete deglycosylation under the conditions tested. Together, these results demonstrate that insect digestion results in partial, rather than complete, deglycosylation of hederagenin-derived saponins.

Glycosylated plant metabolites are typically hydrolyzed by β-glycosidases upon tissue disruption, for example during herbivory or sample processing, when enzymes and substrates come into contact. To test whether such processes could account for the observed transformation, *B. vulgaris* extracts were applied to *Brassica napus* leaf discs and processed without flash freezing in liquid nitrogen, thereby allowing potential plant enzymatic activity in the absence of insects. Under these conditions, no detectable change in saponin composition was observed (Fig. 3). This demonstrates that the observed turnover is not an artefact of plant-derived enzymatic activity during extraction or incubation but instead reflects insect-mediated processes.

### Position-specific sugar removal determines saponin activity and metabolic fate

A key unresolved question is whether deglycosylation of saponins represents detoxification or activation. Importantly, sugar removal can occur at different positions and may therefore generate structurally and functionally distinct products. In our study, hederagenin cellobioside was converted to the corresponding 3-O-monoglucoside, indicating partial removal of sugar moieties at the C3 position. In contrast, the bidesmosidic saponin hederacoside C was not converted to a monodesmosidic form, indicating that removal of the C28-linked sugar chain does not occur under the conditions tested.

The prevailing paradigm assumes that removal of sugar moieties reduces toxicity by increasing hydrophobicity and facilitating excretion. However, previous studies have demonstrated that both hederagenin cellobioside and its corresponding 3-O-monoglucoside are toxic and deterrent to *P. xylostella* (Shinoda et al., 2002; Liu et al., 2019; Augustin et al., 2012). Although these studies differed in larval stage and host plant, they demonstrate that both the di- and monoglycosylated forms are highly bioactive.

Our results further support this conclusion. Enrichment of monoglucosides during insect digestion indicates that partial deglycosylation at C3 does not detoxify these compounds. In parallel, α-hederin, a monodesmosidic saponin lacking a C28 sugar chain, exhibits strong antifeedant activity and toxicity, whereas the corresponding bidesmosidic hederacoside C is inactive. Notably, this difference in activity is not associated with detectable conversion of hederacoside C to α-hederin, indicating that removal of the C28-linked sugar chain does not occur during digestion.

The strong contrast in activity between monodesmosidic and bidesmosidic saponins further highlights the importance of glycosylation pattern. α-Hederin exhibited near-complete antifeedant activity and measurable toxicity, whereas hederacoside C was inactive and not detectably transformed. These findings are consistent with previous reports showing that C3-monodesmosidic hederagenin-derived saponins are more bioactive than bidesmosidic forms (Avato et al., 2006; Tian et al., 2021).

A likely explanation is that glycosylation at C28 alters the physicochemical properties of the molecule, including its amphiphilicity and ability to interact with membrane sterols, thereby reducing its biological activity (Wink et al., 2015; Devishi et al., 2026; Wang et al., 2026). Our results further indicate that glycosylation pattern influences not only intrinsic activity but also susceptibility to metabolic transformation during digestion.

The absence of detectable turnover of hederacoside C was unexpected. The lepidopteran midgut is strongly alkaline, and such conditions might be expected to promote hydrolysis of ester-linked sugar chains at the C28 position. However, no such transformation was detected. This suggests that the C28 ester linkage in hederagenin-derived saponins is more stable under physiological conditions than anticipated and that gut alkalinity alone is insufficient to drive its cleavage. This interpretation is consistent with recent findings showing that these saponins remain stable across a broad pH range, including alkaline conditions (Wang et al., 2026).

Together, these observations demonstrate that the biological consequences of sugar removal depend on both the position and extent of modification. While partial deglycosylation at C3 produces bioactive monoglucosides, the absence of C28 de-esterification prevents conversion of inactive bidesmosidic saponins into more active monodesmosidic forms. In other plant– insect systems, removal of sugar moieties can either neutralize toxicity or generate more potent metabolites depending on compound structure (Vassao et al., 2018; Zhang et al., 2020; Huber et al., 2021). Our results place hederagenin-derived saponins within this broader context and show that insect-mediated turnover is selective rather than indiscriminate, producing a defined set of bioactive defence compounds rather than simply detoxifying them.

### Insect digestion, not plant enzymes, drives saponin transformation during ingestion

Plant tissue is initially ingested under near-neutral conditions before entering the highly alkaline midgut characteristic of lepidopteran larvae (Pentzold et al., 2014). Under these initial conditions, plant-derived β-glucosidases, if present, could potentially remain active and catalyse deglycosylation prior to exposure to alkaline pH. Such two-component defence systems, in which glycosylated precursors are activated upon tissue disruption, are known from other plant–herbivore interactions (Morant et al.,2008).

However, our experiments showed no detectable conversion of *B. vulgaris* saponins in the absence of insects (Fig. 3), indicating that such plant-mediated activation does not occur in this system. This suggests that the observed saponin turnover is primarily driven by insect-associated processes rather than plant enzymes. Instead, the observed turnover appears to be driven primarily by insect-associated processes. This contrasts with systems in which plants tightly control activation of defence compounds and highlights the importance of considering both plant and herbivore contributions to chemical defence dynamics.

### Ecological and evolutionary implications for plant–herbivore interactions

From an ecological and evolutionary perspective, our results suggest that glycosylation plays a dual role in plant–insect interactions by controlling both saponin activity and their transformation during herbivory. In contrast to plant systems in which deglycosylation is tightly regulated and activated upon tissue damage, insect-mediated turnover appears to generate bioactive saponins as a consequence of digestion.

These results raise the possibility that herbivore metabolism may unintentionally activate plant defence compounds, or alternatively, that plants may produce saponin structures that become active following ingestion. In either case, the outcome of plant–herbivore interactions is shaped not only by the compounds produced by the plant but also by how they are transformed within the herbivore.

These findings also have implications for the development of saponins as biological control agents. A detailed understanding of how glycosylation pattern influences both activity and metabolic stability will be important for predicting efficacy against target herbivores. Compounds that remain stable during digestion but retain strong bioactivity, such as monodesmosidic saponins, may represent particularly promising candidates, whereas inactive or transformation-resistant saponins may be less effective.

Thus, mechanistic insight into saponin turnover reveals how glycosylation governs both activity and metabolic fate, providing a foundation for understanding plant–herbivore interactions and for the rational development of saponin-based plant protection strategies.

## ACKNOWLEDGEMENTS

This work was financially supported by Novo Nordisk Foundation (EcoSap NNF20OC0060298 to SB), the China Scholarship Council (CSC, 202108330041 to JS) and Marie Sklodowska-Curie Individual Fellowship (MSCA-IF 752437 to PC).

## AUTHOR CONTRIBUTIONS

JS, PC, and SB designed the research and wrote the paper. JS performed the experiments and analyzed the data.

## REFFERENCES

Agerbirk N, Olsen CE, Bibby BM, Frandsen HO, Brown LD, Nielsen JK, Renwick JAA. 2003. A saponin correlated with variable resistance of Barbarea vulgaris to the diamondback moth Plutella xylostella. Journal of chemical ecology, 29: 1417–1433. 10.1023/A:1024217504445

Akao T, Akao T, Hattori M, Kanaoka M, Yamamoto K, Namba T, Kobashi K. 1991. Hydrolysis of glycyrrhizin to 18β-glycyrrhetyl monoglucuronide by lysosomal β-D-glucuronidase of animal livers. Biochemical pharmacology, 41: 1025–1029. 10.1016/0006-2952(91)90210-V

Augustin JM, Drok S, Shinoda T, Sanmiya K, Nielsen JK, Khakimov B, Olsen CE, Hansen EH, Kuzina V, Ekstrøm CT. 2012. UDP-glycosyltransferases from the UGT73C subfamily in Barbarea vulgaris catalyze sapogenin 3-O-glucosylation in saponin-mediated insect resistance. Plant physiology, 160: 1881–1895. 10.1104/pp.112.202747

Augustin JM, Kuzina V, Andersen SB, Bak S. 2011. Molecular activities, biosynthesis and evolution of triterpenoid saponins. Phytochemistry, 72: 435–457. 10.1016/j.phytochem.2011.01.015

Avato P, Bucci R, Tava A, Vitali C, Rosato A, Bialy Z, Jurzysta M. 2006. Antimicrobial activity of saponins from Medicago sp.: structure-activity relationship. Phytotherapy research, 20: 454–457. 10.1002/ptr.1876

de Jong PW, Breuker CJ, de Vos H, Vermeer KMCA, Oku K, Verbaarschot P, Nielsen JK, Brakefield PM. 2009. Genetic differentiation between resistance phenotypes in the phytophagous flea beetle, Phyllotreta nemorum. Journal of Insect Science, 9: 69.

de Jong PW, Nielsen JK. 2000. Reduction in fitness of flea beetles which are homozygous for an autosomal gene conferring resistance to defences in Barbarea vulgaris. Heredity, 84: 20–28.

Dervishi M, Günther J, Li J, Uzun HD, Hansen HCB, Günther-Pomorski T, Fuglsang AT, Monje V, Bak S. In press. Sterols govern membrane susceptibility to saponin-induced lysis. Proceedings of the National Academy of Sciences of the USA. Preprint available at: 10.1101/2025.07.17.665205

Dervishi M, Schmitt FM, Günther J, Cedergreen N, Bak S. 2026. Structure–activity relationships of triterpenoid saponins across phylogenetically diverse organisms. Journal of Chemical Ecology. 10.1007/s10886-025-01666-3

Divekar PA, Narayana S, Divekar BA, Kumar R, Gadratagi BG, Ray A, Singh AK, Rani V, Singh V, Singh AK. 2022. Plant secondary metabolites as defense tools against herbivores for sustainable crop protection. International journal of molecular sciences, 23: 2690. 10.3390/ijms23052690

Erthmann PØ, Agerbirk N, Bak S. 2018. A tandem array of UDP-glycosyltransferases from the UGT73C subfamily glycosylate sapogenins, forming a spectrum of mono-and bisdesmosidic saponins. Plant Molecular Biology, 97: 37–55. 10.1007/s11103-018-0723-z

Gao GC, Lu ZX, Tao SH, Zhang S, Wang FZ. 2011. Triterpenoid saponins with antifeedant activities from stem bark of Catunaregam spinosa (Rubiaceae) against Plutella xylostella (Plutellidae). Carbohydrate Research, 346: 2200–2205. 10.1016/j.carres.2011.07.022

Khakimov B, Tseng LH, Godejohann M, Bak S, Engelsen SB. 2016. Screening for triterpenoid saponins in plants using hyphenated analytical platforms. Molecules, 21: 1614. 10.3390/molecules21121614

Kuzina V, Ekstrøm CT, Andersen SB, Nielsen JK, Olsen CE, Bak S. 2009. Identification of defense compounds in Barbarea vulgaris against the herbivore Phyllotreta nemorum by an ecometabolomic approach. Plant physiology, 151: 1977–1990. 10.1104/pp.109.136952

Liu Q, Khakimov B, Cárdenas PD, Cozzi F, Olsen CE, Jensen KR, Hauser TP, Bak S. 2019. The cytochrome P450 CYP72A552 is key to production of hederagenin-based saponins that mediate plant defense against herbivores. New Phytologist, 222: 1599–1609. 10.1111/nph.15689Digital Object Identifier (DOI)

Morant AV, Jørgensen K, Jørgensen C, Paquette SM, Sánchez-Pérez R, Møller BL, Bak S. 2008. β-Glucosidases as detonators of plant chemical defense. Phytochemistry, 69: 1795–1813. 10.1016/j.phytochem.2008.03.006

Moses T, Papadopoulou KK, Osbourn A. 2014. Metabolic and functional diversity of saponins, biosynthetic intermediates and semi-synthetic derivatives. Critical reviews in biochemistry and molecular biology, 49: 439–462. 10.3109/10409238.2014.953628

Fürstenberg-Hägg J, Zagrobelny M, Jørgensen K, Vogel H, Møller BL, Bak S. 2014. Chemical defense balanced by sequestration and de novo biosynthesis in a lepidopteran specialist. PLoS ONE, 9: e108745. 10.1371/journal.pone.0108745

Nielsen JK, Nagao T, Okabe H, Shinoda T. 2010. Resistance in the plant, Barbarea vulgaris, and counter-adaptations in flea beetles mediated by saponins. Journal of Chemical Ecology, 36: 277–285. 10.1007/s10886-010-9758-6

Pentzold S, Zagrobelny M, Rook F, Bak S. 2014. How insects overcome two-component plant chemical defence: plant β-glucosidases as the main target for herbivore adaptation. Biological Reviews, 89: 531–551. 10.1111/brv.12066

Qiao X, Li Q, Yin H, Qi K, Li L, Wang R, Zhang S, Paterson AH. 2019. Gene duplication and evolution in recurring polyploidization–diploidization cycles in plants. Genome biology, 20: 38. 10.1186/s13059-019-1650-2

Shen J, Günther J, Kjeldgaard-Nintemann S, Cárdenas PD, Bak S. In press. Metabolic engineering reveals LUP5 as a determinant of saponin composition and insect resistance in Barbarea vulgaris. Plant Physiology. Preprint available at: 10.1101/2025.08.04.665341

Shinoda T, Nagao T, Nakayama M, Serizawa H, Koshioka M, Okabe H, Kawai A. 2002. Identification of a triterpenoid saponin from a crucifer, Barbarea vulgaris, as a feeding deterrent to the diamondback moth, Plutella xylostella. Journal of Chemical Ecology, 28: 587–599. 10.1023/A:1014500330510

Tian X, Li Y, Hao N, Su X, Du J, Hu J, Tian X. 2021. The antifeedant, insecticidal and insect growth inhibitory activities of triterpenoid saponins from Clematis aethusifolia Turcz against Plutella xylostella (L.). Pest Management Science, 77: 455–463. 10.1002/ps.6038

Vats TK, Rawal V, Mullick S, Devi MR, Singh P, Singh AK. 2019. Bioactivity of Ageratum conyzoides (L.)(Asteraceae) on feeding and oviposition behaviour of diamondback moth Plutella xylostella (L.)(Lepidoptera: Plutellidae). International Journal of Tropical Insect Science, 39: 311–318. 10.1007/s42690-019-00042-5

Wang C, Kirkensgaard JJK, Risbo J, Dervishi M, Nahapetyan G, Hassenkam T, Rami M, Bak S, Hansen HCB. 2026. Glycosylation pattern controls solubility, micellization, and aggregation of structurally defined ivy-derived triterpenoid saponins. JCIS Open, 22: 100177. 10.1016/j.jciso.2026.100177

